# Pharmacological Inhibition of FKBP51 Mitigates Early Life Adversity-Induced Social Deficits

**DOI:** 10.64898/2025.12.13.694102

**Authors:** Joeri Bordes, Xiuqi Ji, Serena Gasperoni, Choham Sudre-Chinsky, Daniela Harbich, Cornelia Flachskamm, Paula Fontanet, Sowmya Narayan, Manfred Uhr, Christian Namendorf, Alon Chen, Felix Hausch, Juan Pablo Lopez, Mathias V. Schmidt

## Abstract

Early life adversity (ELA) is a major risk factor for psychiatric disorders, but targeted preventative strategies are lacking due to poor mechanistic insight. The FKBP51 protein, a co-chaperone of the glucocorticoid receptor, is a key mediator of stress vulnerability. We tested if pharmacological inhibition of FKBP51 with the selective inhibitor SAFit2 prevents the long-term consequences of ELA. Mice exposed to ELA exhibited persistent deficits in social behavior, manifesting as social subordination in adolescence and adulthood. Early-life SAFit2 treatment fully rescued these ELA-induced behavioral impairments. Transcriptional profiling across six stress-relevant brain regions revealed that SAFit2 normalized ELA-driven gene expression changes, particularly in the medial prefrontal cortex and nucleus accumbens. Functional analysis showed the rescue converged on immunoregulatory and neuroactive ligand-receptor signaling pathways. Our findings establish FKBP51 as a critical pharmacological target for reversing the lasting impact of early life adversity on brain function, offering a path toward preventative treatment for ELA-related psychopathology.

## Main

Early life adversity (ELA), including childhood trauma and neglect, is a significant societal problem with profound and lasting consequences for mental health. Epidemiological studies consistently show that ELA is one of the most powerful risk factors associated with the development of major psychiatric disorders, including Major Depressive Disorder (MDD), anxiety, and Post-Traumatic Stress Disorder (PTSD), among others^1–3^. While a direct association is clear, the underlying molecular and neurobiological mechanisms by which early adversity becomes biologically embedded and contributes to lifelong vulnerability remain poorly understood. This lack of mechanistic insight has hampered the development of targeted, preventative interventions that could be implemented during or shortly after periods of early trauma.

A growing body of evidence highlights FK506-binding protein 5 (FKBP5) gene as a key mediator of gene-environment interactions underlying the long-term impact of ELA on psychiatric vulnerability. Notably, the rs1360780 polymorphism in the FKBP5 gene has been consistently associated with an increased risk for MDD, PTSD, and other stress-related disorders, especially among individuals exposed to childhood trauma^4–9^. The protein encoded by this gene, FKBP51, is a key co-chaperone of the glucocorticoid receptor (GR), modulating its sensitivity to stress hormones. This interaction reduces GR sensitivity to glucocorticoids, weakening the negative feedback regulation of the hypothalamic-pituitary-adrenal (HPA) axis^10,11^. The resulting dysregulation of the HPA axis, characterized by prolonged elevation of stress hormones, is a hallmark of many stress-related psychiatric disorders. Supporting this, post-mortem studies have shown increased FKBP51 expression in the brains of individuals with MDD and PTSD, indicating long-lasting molecular alterations^12,13^. In parallel, Fkbp5 knockout mouse models reveal a strong causal role for this gene in orchestrating individual differences in stress resilience and susceptibility, reinforcing its relevance in the pathophysiology of stress-related disorders^14–18^. Recent preclinical studies have begun to map the brain region-specific transcriptional consequences of ELA, revealing how early adversity becomes biologically embedded through persistent gene expression changes across key circuits. For example, our prior work demonstrated that ELA disrupts social hierarchy and induces cell-type-specific transcriptomic alterations in the ventral hippocampus, contributing to excitation/inhibition imbalance and behavioral dysfunction^19^. This expanding knowledge base offers a roadmap for identifying molecular targets and tracking the reversal of ELA-induced pathology.

The convergence of genetic, molecular, and behavioral evidence positions FKBP51 as a compelling therapeutic target for mitigating the long-lasting impact of early trauma. Until recently, a key limitation has been the absence of a selective, brain-penetrant small molecule capable of pharmacologically inhibiting FKBP51 in vivo. The recent development of SAFit2, a highly selective and potent FKBP51 inhibitor, has provided a critical pharmacological tool to directly evaluate the therapeutic potential of FKBP51 inhibition^20–24^. SAFit2 exhibits excellent brain permeability and effectively modulates GR function in preclinical models, opening new avenues for therapeutic exploration.

Alongside these molecular insights, behavioral neuroscience has made significant strides, particularly in the domain of social behavior. Social deficits are a hallmark of numerous psychiatric disorders and represent a persistent consequence of ELA. While conventional behavioral assays have provided fundamental insights, recent advances in deep behavioral phenotyping and computer vision-based tracking now allow for high-resolution, unbiased quantification of complex social interactions in vivo^25–28^. These sophisticated approaches deconstruct global social impairments into discrete, quantifiable components, offering a detailed view of how ELA reshapes social behavior and enabling more sensitive evaluation for the efficacy of therapeutic interventions^29^.

In this study, we leveraged the selective FKBP51 inhibitor SAFit2 to test whether pharmacological intervention can abolish the long-term neurobehavioral and transcriptional consequences of ELA. Mice were exposed to a well-established model of ELA and treated with either SAFit2 or vehicle control. Through automated deep behavioral phenotyping, we demonstrate that SAFit2 treatment successfully rescues ELA-induced deficits in social behavior, in both adolescence and young adulthood. Furthermore, we performed bulk RNA sequencing across multiple stress-relevant brain regions and demonstrate that SAFit2 normalizes ELA-induced transcriptional changes. These data uncover a complex regulatory landscape, with some genes showing cross-regional rescue, while others exhibiting region-specific responses to SAFit2. Together, our findings support FKBP51 as a promising therapeutic target for reversing the lasting impact of early life adversity and shed light on the transcriptional mechanisms that mediate this rescue.

## Results

### The role of SAFit2 in shaping physiological hallmarks of early life adversity

To determine whether SAFit2 modulates the physiological effects of early life adversity (ELA), dams received a chronic injection of SAFit2 or vehicle starting at postnatal day 2 (P02), coinciding with the onset of the ELA paradigm (**Figure 1A**). This design enabled comparison of baseline conditions (Ctrl_Veh), potential stress effects in the absence of SAFit2 (ELA_Veh), and the ability of SAFit2 to counteract ELA-induced alterations (ELA_SAFit2). The transfer of SAFit2 to the offspring was verified by detecting stable concentrations of the compound in pup plasma and stomach contents at P04, P07, and P09 (**Figure 1B**).

**Figure 1.**
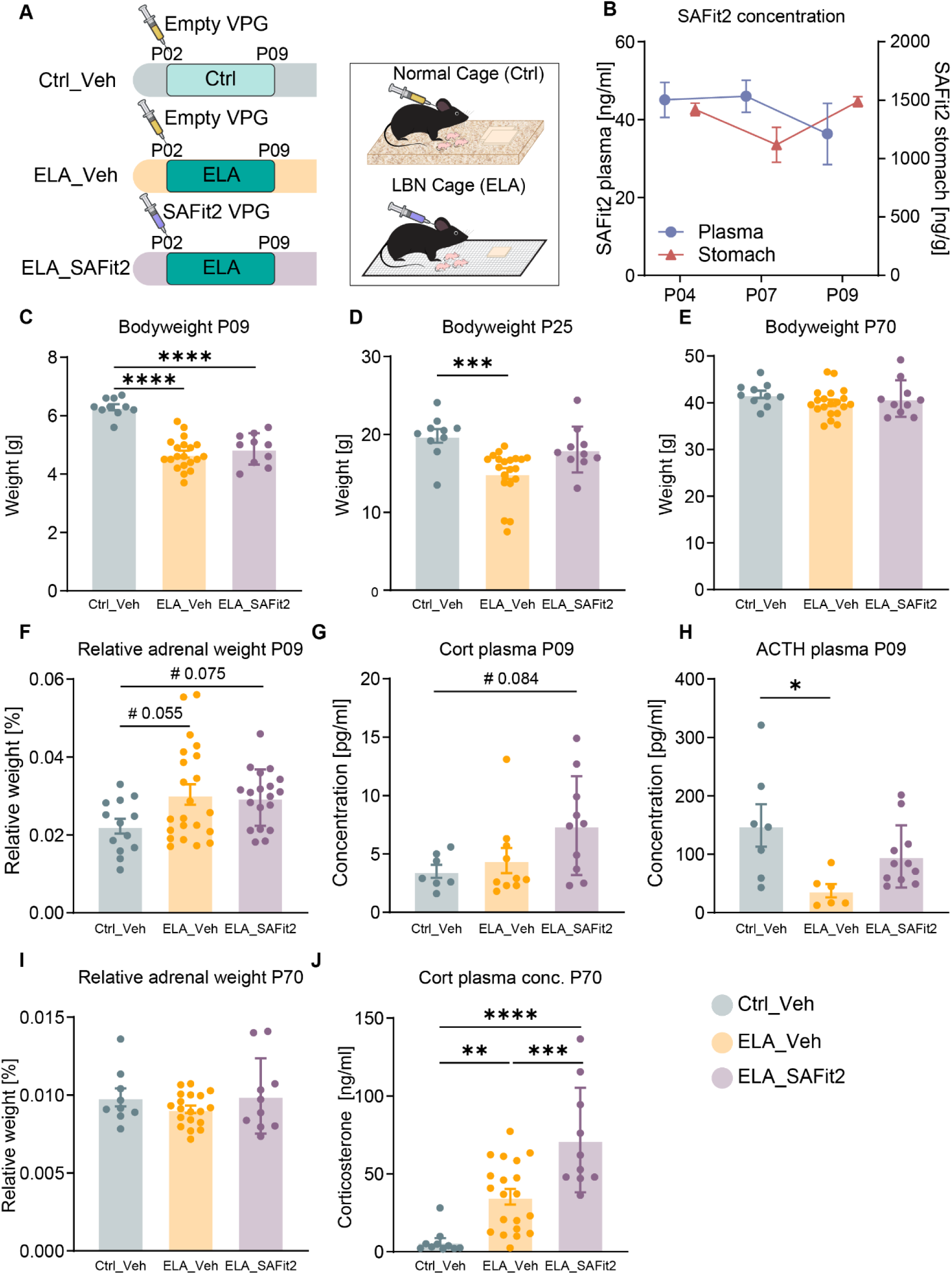
The role of SAFit2 in shaping physiological hallmarks of early life stress. **A)**. Experimental timeline of the nonstressed and stressed conditions with the pharmacological intervention using SAFit2. **B)** The SAFit2 concentration in blood plasma and stomach in the pups during stress exposure. **C)** Body weight at P09. One-way ANOVA showed a main effect, F (2,37) = 38.49, p < 0.0001. Follow-up Holm-Šídák tests showed lower body weight in both ELA groups vs Ctrl_veh (p-values<0.0001). **D)** Body weight at P25. Kruskal-Wallis test showed a main effect, H (2) = 16.31, p = 0.0003. Follow-up Dunn’s tests showed lower body weight in ELA_Veh vs Ctrl_veh (p=0.0002), but not in ELA_SAFit2 vs Ctrl_veh (p=0.3247). **E)** Adult weight. One-way ANOVA showed no effect, F (2,37) = 1.099, p = 0.3439. **F)** Relative adrenal weight at P09. One-way ANOVA showed a main effect, F(2,51) = 3.279, p = 0.0457. Follow-up Holm-Šídák tests detected no pairwise differences for ELA_Veh vs ELA_SAFit2: p=0.7881, although a trend was observed for Ctrl_veh vs ELA_Veh: p = 0.0550 and Ctrl_veh vs ELA_SAFit2: p=0.0753 **G)** Baseline corticosterone plasma levels at P09. One-way ANOVA showed a trend toward a group effect, F(2,24) = 3.208, p = 0.0583. Follow-up Holm-Šídák tests indicated a trend for Ctrl_veh vs ELA_SAFit2 (p = 0.0844), but not for Ctrl_veh vs ELA_Veh (p = 0.5870) or ELA_Veh vs ELA_SAFit2 (p = 0.1212). **H)** Baseline ACTH plasma levels at P09. One-way ANOVA showed a main effect, F(2,21) = 4.829, p = 0.0188. Follow-up Holm–Šídák tests showed higher levels in Ctrl_veh vs ELA_Veh (p = 0.0159), with no differences for Ctrl_veh vs ELA_SAFit2 (p = 0.1673) or ELA_Veh vs ELA_SAFit2 (p = 0.1673). **I)** Relative adrenal weight in adulthood. One-way ANOVA showed no effect, F(2,35) = 1.155, p = 0.3268. **J)** Baseline corticosterone plasma levels in adulthood. One-way ANOVA showed a main effect, F(2,37) = 19.81, p < 0.0001. Follow-up Holm–Šídák tests showed higher levels in both ELA groups vs Ctrl_veh (Ctrl_veh vs ELA_Veh: p = 0.0027; Ctrl_veh vs ELA_SAFit2: p < 0.0001) and in ELA_SAFit2 vs ELA_Veh (p = 0.0005). Sample size: panels B-E, I-J: n=10 for Ctrl_veh (1 excluded in panel I), n=20 for ELA_Veh (1 excluded in panel I), n=10 for ELA_SAFit2. Panel F: n=13 for Ctrl_veh, n=22 for ELA_Veh, n=19 for ELA_SAFit2. Panel G: n=7 for Ctrl_veh, n=10 for ELA_Veh, n=10 for ELA_SAFit2. Panel H: n=7 for Ctrl_veh, n=6 for ELA_Veh, n=11 for ELA_SAFit2.

A well-established consequence of limited bedding and nesting (LBN) exposure is reduced body weight at P09^19,30,31^. In the present study, body weight was significantly reduced at P09 after ELA, independent of SAFit2 or vehicle treatment, confirming successful induction of chronic stress that was not prevented by SAFit2 at this early stage (**Figure 1C**). At P25, the stress-induced reduction in body weight persisted in ELA vehicle animals but not in ELA_SAFit2 animals, indicating a faster recovery to baseline body weight (**Figure 1D**). By adulthood (P70), no differences in body weight were detected among Ctrl_Veh, ELA_Veh, and ELA_SAFit2 groups, demonstrating complete recovery in both stressed groups (**Figure 1E**).

Next, classical hallmarks of HPA-axis dysregulation caused by chronic stress exposure were investigated via analysis of the relative adrenal weight, basal adrenocorticotropic hormone (ACTH), and basal corticosterone directly after stress (P09) and into young adulthood (P70). The relative adrenal weight at P09 did not show a significant difference across conditions, although both the ELA groups showed a trend toward higher adrenal weight relative to Ctrl_Veh (**Figure 1F**). Furthermore, baseline corticosterone at P09 showed a trend toward higher levels in ELA_SAFit2 vs Ctrl_Veh (**Figure 1G**), while baseline ACTH at P09 was significantly higher in Ctrl_Veh vs ELA_Veh, with no differences between Ctrl_Veh and ELA_SAFit2 or between ELA_Veh and ELA_SAFit2 (**Figure 1H**), indicating condition-specific regulation of the HPA axis in response to early life adversity and SAFit2 treatment. In adulthood, relative adrenal weight did not differ among conditions (**Figure 1I**). However, adult baseline corticosterone showed significantly higher levels in both ELA groups vs Ctrl_Veh, and also significantly elevated levels in ELA_SAFit2 vs ELA_Veh.

### ELA-induced changes in social behaviors are persistent and rescued by SAFit2

To investigate the long-term impact of ELA and the potential rescuing effect of SAFit2 intervention on social behaviors, we formed ten mix-condition groups, each containing four male mice, one Ctrl_Veh, two ELA_Veh and one ELA_SAFit2 animal. Males were specifically selected due to the previously reported stronger ELA-induced effects on social behavior^19^. These groups were assessed using the social box (SB) system, a semi-naturalistic, enriched living environment that enables automated, continuous tracking of freely interacting animals. Behavioral recordings were conducted during two key developmental stages: adolescence (postnatal day 30, P30) and young adulthood (P61) (**Figure 2A**). The SB setup allowed for high-resolution, longitudinal analysis of complex social interactions within a group context.

**Figure 2.**
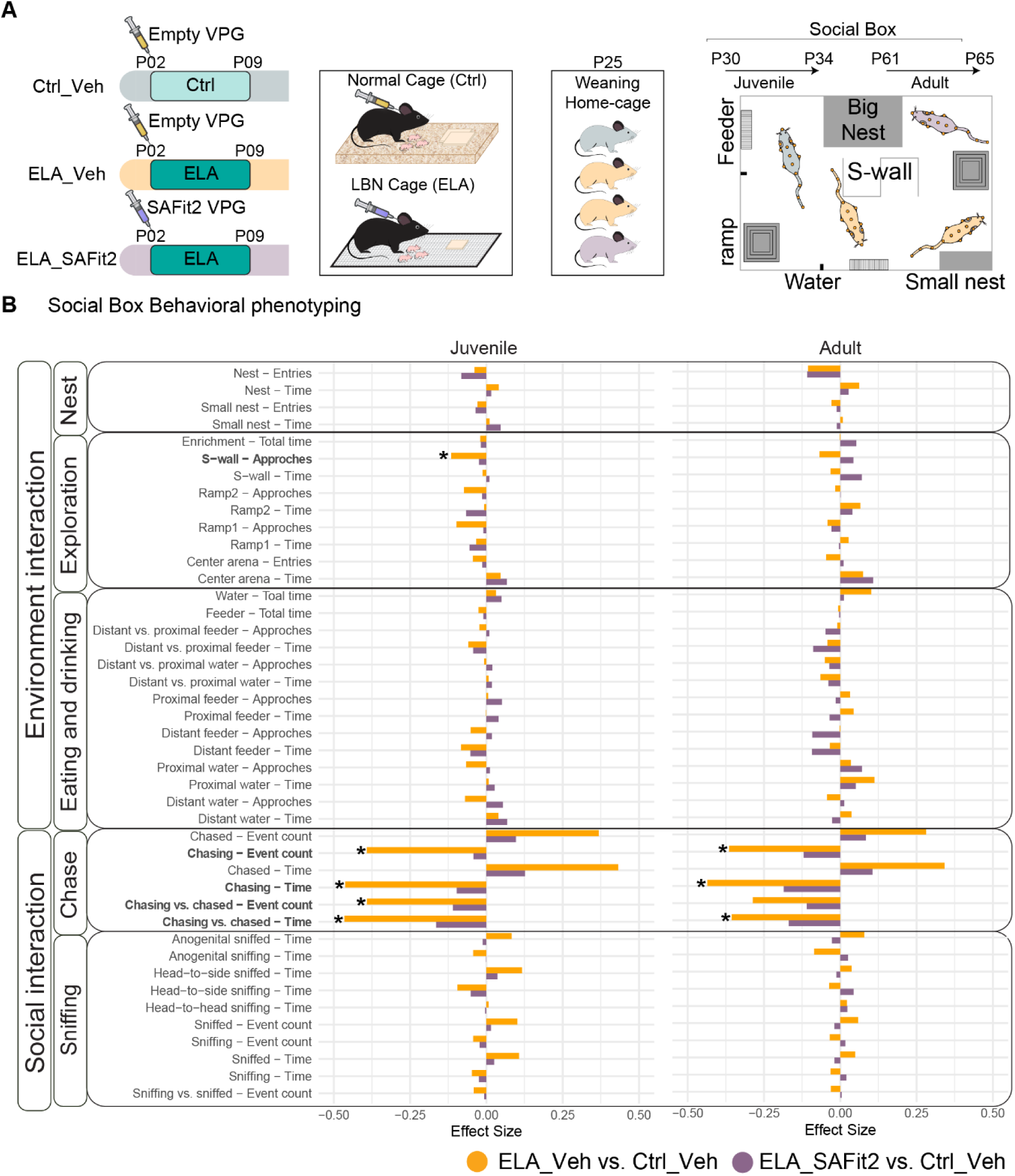
SAFit2 rescues ELA-induced changes in social behaviors. **A)**. Schematic representation of the experimental timeline from birth to behavioral testing showing the three conditions: control treated with vehicle (Ctrl_Veh), ELA exposed treated with vehicle (ELA_Veh), and ELA exposed treated with SAFit2 (ELA_SAFit2). Social behavior was assessed during adolescence (P30) and early adulthood (P61) using the Social Box (SB), an enriched semi-naturalistic environment enabling continuous behavioral tracking. **B).** Summary of behaviors at P30 and P61, comparing ELA_Veh vs. Ctrl_Veh and ELA_SAFit2 vs. Ctrl_Veh using a linear mixed-effect model. FDR adjustment was applied on p-values. At the juvenile stage, ELA_Veh showed significantly lower total chasing time (β = -0.463, SE = 0.177, *p = 0.032*), event counts (β = -0.392, SE = 0.156, *p = 0.040*), the chase-to-being-chased ratio in time (β = -0.465, SE = 0.163, *p = 0.019*) and event count (β = -0.391, SE = 0.143, *p = 0.025*), and the number of approaches to the s-wall (β = -0.115, SE = 0.047, *p = 0.045*); in adulthood, ELA_Veh showed significantly reduced chasing time (β = -0.435, SE = 0.145, *p = 0.011*), event counts (β = -0.364, SE = 0.125, *p = 0.014*), and the chase-to-being chased ratio in time (β = -0.356, SE = 0.137, *p = 0.029*). Sample size: n=10 for Ctrl_veh, n=20 for ELA_Veh, n=10 for ELA_SAFit2.

We obtained a total of 43 ethologically relevant behavioral readouts across four days (comprising four active and three inactive phases) during both juvenile (P30) and adult (P61) stages. The behaviors were categorized into environmental interactions (nest, exploration, eating, and drinking) and social interactions (chasing, sniffing) (**Figure 2B**). Linear mixed-effects modeling revealed that ELA_Veh animals exhibited significant reductions in chase-related behaviors at both developmental stages compared to Ctrl_Veh animals (**Figure 2B**). At the juvenile stage, total chasing time and event counts, the chase-to-being-chased ratio, and the number of approaches to the s-wall were all significantly lower in ELA_Veh (**Figure 2B**). Similarly, in adulthood, ELA_Veh showed a significantly reduced chase-to-being-chased ratio compared to Ctrl_Veh (**Figure 2B**). In contrast, ELA_SAFit2 mice did not differ significantly from Ctrl_Veh in any of the measured behaviors at either stage (**Figure 2B**). These findings suggest that ELA leads to persistent deficits in social behavior, which are effectively mitigated by SAFit2 treatment.

Finally, to investigate the temporal dynamics of behavioral alterations, readouts from the active phases were segmented into 2-hour intervals and compared across Ctrl_Veh, ELA_Veh and ELA_SAFit2 conditions. For behavioral features that showed significant differences between ELA_Veh and Ctrl_Veh, these differences were consistently observed across all 2-hour time windows during the active phases (**Supplementary Figure 1**), indicating that the group-specific behavioral patterns were stable and persistent throughout the active period. These results underscore the robustness of ELA-induced social deficits and highlight SAFit2 as a promising intervention capable of restoring stable behavioral dynamics across developmental stages and time.

### ELA-induced social subordination is abolished by FKBP51 inhibition via SAFit2 treatment

Social hierarchy provides a powerful representation of the social dynamics in group-housed mice. To assess the impact of ELA and SAFit2 treatment on social dominance, we applied the well-established David’s score method^32,33^. David’s score is a widely used metric for quantifying social dominance based on agonistic interactions. It calculates the proportion of wins (chasing) and losses (being chased) relative to the total number of such interactions within a group, providing a continuous measure of individual dominance status derived from observed social behavior. At both adolescent and adult stages, ELA_Veh mice exhibited significantly lower David’s scores compared to Ctrl_Veh mice, while ELA_SAFit2 animals did not differ significantly from controls (**Figure 3A**). This pattern was consistently observed across all four days of behavioral assessment (**Figure 3A**). The distribution of social ranks within each group also revealed significant differences across conditions (**Figure 3B-C**). ELA_Veh mice were disproportionately represented in lower social ranks (gamma and delta) relative to Ctrl_Veh and ELA_SAFit2 animals (**Figure 3B**). Conversely, fewer ELA_Veh mice occupied higher social ranks (alpha and beta), whereas ELA_SAFit2 mice showed a rank distribution comparable to Ctrl_Veh mice, with a greater proportion in dominant positions (**Figure 3C**). Together, these findings not only reinforce the link between early life adversity and social subordination, but also demonstrate that FKBP51 inhibition via SAFit2 treatment can effectively restore normative social hierarchy dynamics, offering a promising avenue for therapeutic intervention.

**Figure 3.**
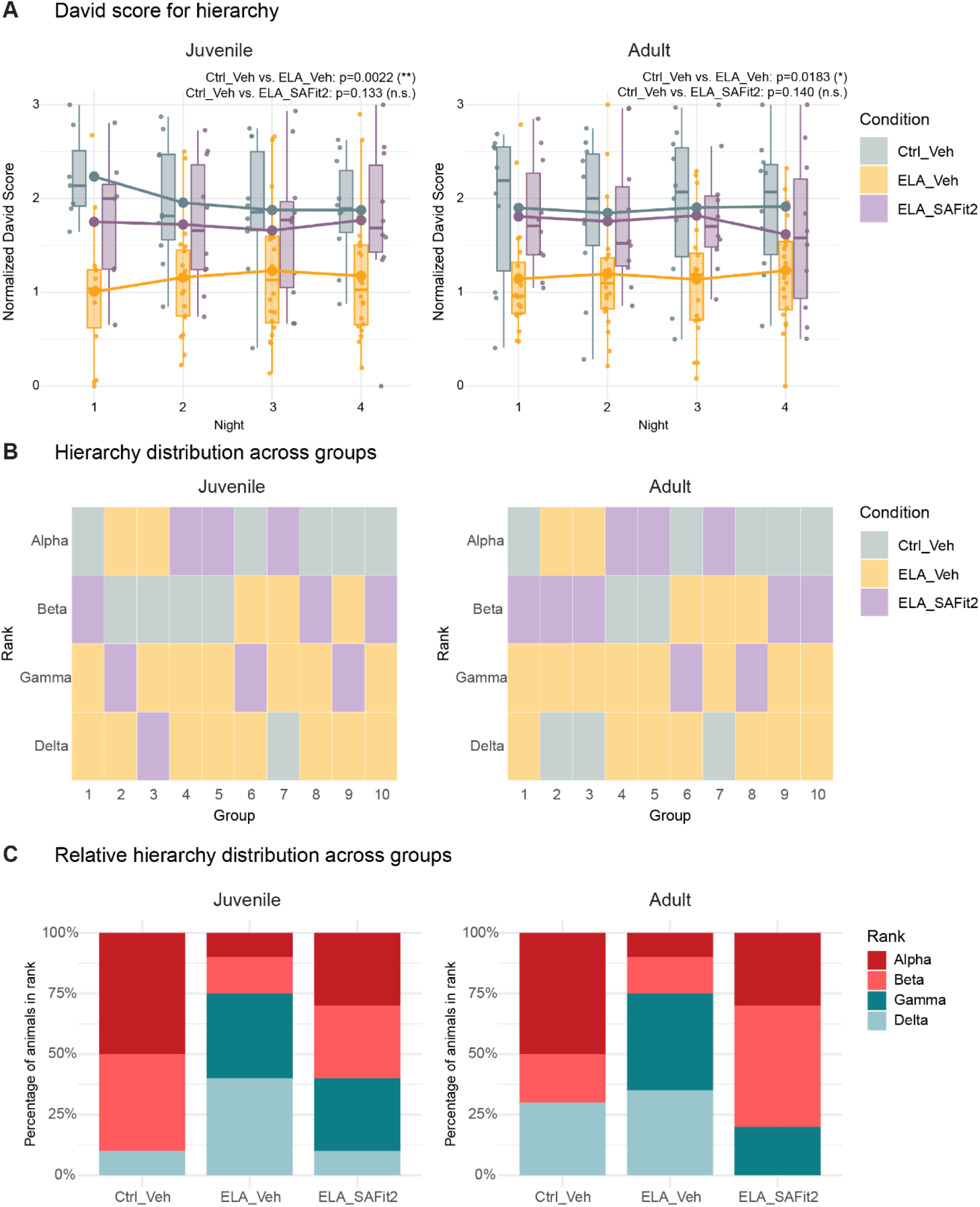
SAFit2 normalized ELA-induced social subordination. **A).** David’s score of juvenile and adult animals. ELA_Veh mice showed significantly lower scores compared to Ctrl_Veh (*juvenile: F (1,95) = 9.91, p = 0.002; adult: F (1,104) = 5.74, p = 0.018*) while ELA_SAFit2 showed no significant difference (*juvenile: F (1,58) = 2.32, p = 0.133; adult: F (1,64) =2.24, p = 0.140*). Two-way repeated-measures ANOVA. **B).** Hierarchy distribution at juvenile and adult stages based on the cumulative David’s score over 4 days. The hierarchy order from the highest ranking to lowest is alpha, beta, gamma, delta. ELA_Veh showed significantly different rank distribution compared to Ctrl_Veh (*juvenile: χ^2^ (3, N = 30) = 11.86, p = 0.008; adult: χ^2^ (3, N = 30) = 8.72, p = 0.033*), while ELA_SAFit2 showed no significant differences (*juvenile: χ^2^ (3, N = 20) = 3.64, p = 0.303; adult: χ^2^ (3, N = 20) = 6.79, p = 0.079*). Yate’s corrected chi-square test. **C).** Relative hierarchy distribution within Ctrl_Veh, ELA_Veh and ELA_SAFit2 at juvenile and adult stages. ELA_Veh has a lower percentage of dominant ranks (alpha, beta) than Ctrl_Veh (*juvenile: p = 0.001, odds ratio = 23.56, 95% CI [2.35, 1248.37]; adult: p = 0.045, odds ratio = 6.48, 95% CI [1.01, 55.16]*), while ELA_SAFit2 does not differ from Ctrl_Veh (*juvenile: p = 0.303, odds ratio = 5.49, 95% CI [0.41, 326.83]; adult: p = 1.00, odds ratio = 0.60, 95% CI [0.04, 6.94].*). Fisher’s exact test. Sample size: n=10 for Ctrl_veh, n=20 for ELA_Veh, n=10 for ELA_SAFit2.

### SAFit2 restores ELA-disrupted transcriptional changes in multiple stress-related brain regions

To investigate how ELA and the pharmacological modulation via SAFit2 shape long-term gene expression, we performed bulk RNA sequencing on samples from six brain regions of adult offspring of the three conditions: Ctrl_Veh, ELA_Veh and ELA_SAFit2 (n=7 per condition). The selected regions are key areas in neural circuits regulating stress, emotion and social behavior. Specifically, the medial prefrontal cortex (mPFC) regulates top-down control of stress responses and social interactions, the anterior cingulate cortex (ACC) is central to emotional evaluation and stress reactivity, the dorsomedial thalamus (DMT) serves as an integrative hub linking prefrontal and limbic structures, the nucleus accumbens (Nacc) mediates motivation and reward sensitivity under stress, the basolateral amygdala (BLA) is critical for emotional memory and fear learning, and the ventral hippocampus (vHipp) contributes to stress regulation and contextual processing.

We performed differential gene expression analysis with pairwise comparisons between Ctrl_Veh, ELA_Veh and ELA_SAFit2. Only a limited number of genes were identified as differentially expressed with statistical significance (adjusted p<0.05), likely reflecting the subtle and regionally dispersed transcriptional changes that emerged in adulthood following ELA and SAFit2 treatment. Nevertheless, certain brain regions exhibited more pronounced shifts, with genes showing consistent fold-change trends and lower p-values. To further explore the transcriptomic data and capture biologically meaningful regulation, we filtered for genes with absolute Log2 fold change greater than 0.5 in at least one pairwise comparison, capturing those with robust and condition-specific expression changes. The analysis also revealed distinct sets of genes whose expression was altered by ELA and, in some cases, restored by SAFit2 treatment. To systematically identify such gene expression trajectories, we devised and applied a pattern scoring approach to detect genes exhibiting changes consistent with ELA-induced dysregulation and SAFit2-mediated rescue. A higher pattern score indicates a greater likelihood that a gene follows this regulatory profile (**Figure 4A**). Pattern scores were calculated for all genes meeting the fold-change threshold across the six regions. When comparing across brain regions, we observed variation in both the number of dysregulated genes and the distribution of pattern scores (**Figure 4B**). Notably, the mPFC, Nacc and BLA showed more dynamic transcriptomic responses and a higher degree of gene patterning, while the remaining regions exhibited more modest shifts (**Figure 4B**). We also identified region-specific regulation patterns among the top-scoring genes (**Figure 4C**). In the mPFC, the top 30 highest scoring genes were predominantly downregulated by ELA and upregulated by SAFit2 treatment, whereas in the Nacc, the opposite pattern was observed, with an upregulation by ELA and downregulation by SAFit2 treatment (**Figure 4C**).

**Figure 4.**
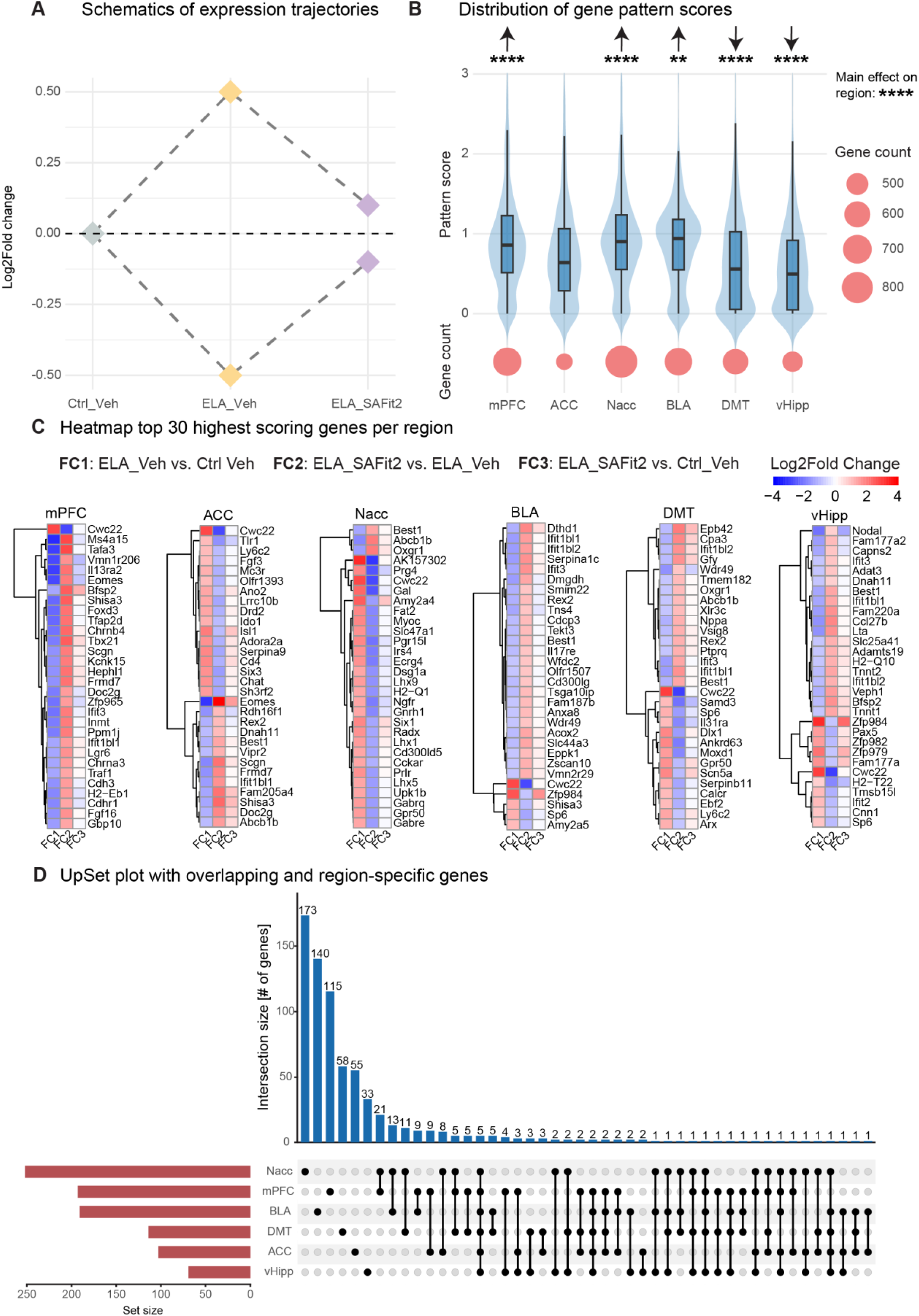
Distinct and region-specific regulation patterns by ELA and SAFit2. **A).** Schematic illustration of the expression trajectories following ELA and SAFit2 treatment identified using the pattern scoring approach. **B).** Distribution of gene pattern scores and the number of dysregulated genes (|Log2FC| > 0.5) across six brain regions: medial prefrontal cortex (mPFC), anterior cingulate cortex (ACC), nucleus accumbens (Nacc), basolateral amygdala (BLA) dorsomedial thalamus (DMT), and ventral hippocampus (vHipp). The pattern score distribution differs significantly across regions (*F (5,3677) = 33.08, p = 2^e-^*^16^). Pattern scores were significantly higher in the mPFC (*estimate =0.17, p < 0.0001*), NAcc (*estimate = 0.16, p < 0.0001*), and BLA (*estimate = 0.07, p = 0.005*). In contrast, DMT (*estimate = –0.15, p < 0.0001*) and vHipp (*estimate = –0.21, p < 0.0001*) showed significantly lower pattern scores compared with the overall mean, while ACC did not differ significantly (*estimate = –0.03, p = 0.23*). All p-values were corrected using FDR. One way ANOVA and post hoc estimated marginal means with contrast effects. **C).** Heatmap visualization of the log2 fold change of the top 30 highest scoring genes in each region. The three columns in each heatmap represent the three differential expression comparisons: FC1 = ELA_Veh vs. Ctrl Veh, FC2 = ELA_SAFit2 vs. ELA_Veh, FC3 = ELA_SAFit2 vs. Ctrl_Veh. **D).** UpSet plot showing the number of overlapping and region-specific genes with high pattern score (>1.15) across the six regions. Sample size: n=7 for Ctrl_veh, n=7 for ELA_Veh, n=7 for ELA_SAFit2.

To examine the convergence and divergence of genes with high pattern scores, we identified both region-specific and cross-regional gene sets (**Figure 4D**). While most genes displaying the pattern were unique to individual regions, a subset of genes followed the ELA-SAFit2 regulation trajectory across multiple brain areas (**Figure 4D**), including *Fkbp5* (**Figure 5A**). Among this subset, *Abcb1b*, a transporter gene implicated in blood-brain barrier regulation and stress responsivity, was downregulated by ELA and restored by SAFit2 (**Figure 5B**). Similarly, *Best1*, associated with astrocytic signaling and synaptic plasticity, showed reduced expression in ELA_Veh, which was restored in the ELA_SAFit2 condition (**Figure 5C**). Finally, the expression of *Tlr1*, a key component of the innate immune signaling pathway, was upregulated by ELA in five out of six regions, and attenuated by SAFit2 treatment (**Figure 5D**). These consistent expression trajectories across multiple regions suggest a shared molecular mechanism underlying ELA-induced vulnerability and its reversal through FKBP51 inhibition by SAFit2 treatment.

**Figure 5.**
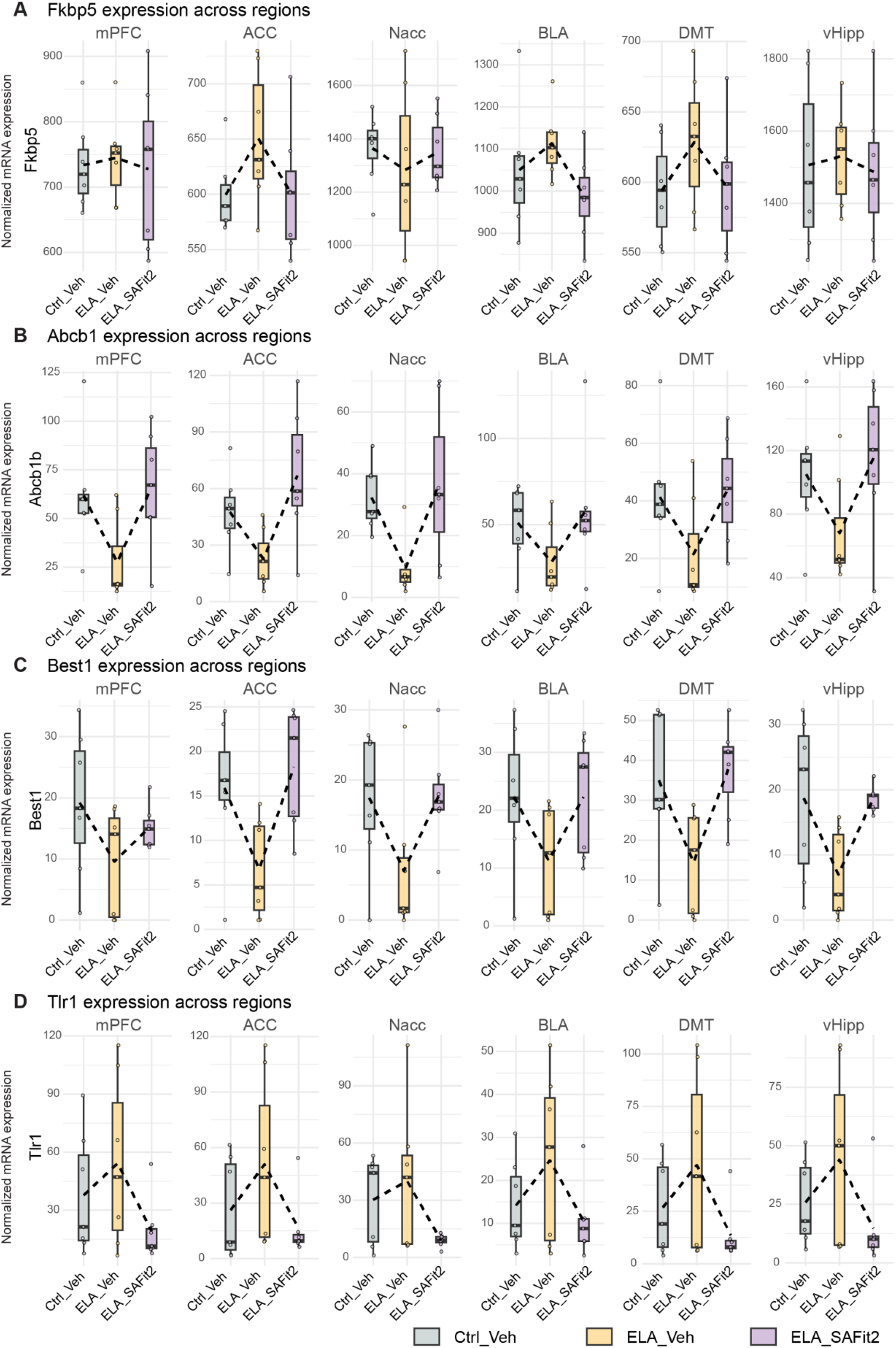
Genes of interest show consistent ELA-SAFit2 expression trajectories across brain regions. **A).** *Fkbp5* expression increased following ELA in ACC, DMT, BLA, vHipp and was normalized by SAFit2. **B).** ELA-induced downregulation of *Abcb1b* in six brain regions restored by SAFit2. **C).** ELA-induced downregulation of *Best1* in six brain regions restored by SAFit2. **D).** ELA-induced upregulation of *Tlr1* in six brain regions restored by SAFit2. Expression across genes is displayed in normalized mRNA expression values. Sample size: n=7 for Ctrl_veh, n=7 for ELA_Veh, n=7 for ELA_SAFit2.

### ELA-SAFit2-regulated genes in mPFC and Nacc reveal distinct functional enrichment in immune and neuroactive signaling pathways

To uncover the biological significance of genes following the ELA-SAFit2 regulation trajectory, we performed functional clustering using the Metascape gene annotation and functional enrichment tool^34^, focusing on two brain regions with the most pronounced transcriptional patterning, the medial prefrontal cortex (mPFC) and nucleus accumbens (Nacc) (**Supplementary Figure 2**). These regions were selected based on their distinct gene expression responses to ELA and SAFit2 treatment, as well as their central roles in stress regulation and social behavior. Genes with elevated pattern scores, indicating ELA-induced dysregulation and SAFit2-mediated rescue, were categorized into three groups: mPFC-specific, Nacc-specific, and overlapping (present in both regions). Functional clusters were then identified and visualized as network maps to reveal shared and region-specific biological pathways (**Supplementary Figure 2A**). Genes overlapping between mPFC and Nacc were significantly enriched in immunoregulatory interactions, suggesting a common inflammatory or immune-related mechanism underlying ELA-induced vulnerability and its reversal. In contrast, Nacc-specific genes were predominantly associated with neuroactive ligand-receptor signaling, pointing to regionally distinct modulation of neurotransmission and reward-related processes (**Supplementary Figure 2B**). These findings highlight that while SAFit2 exerts a broad rescuing effect on ELA-induced transcriptional changes, the underlying biological pathways are regionally specialized. The convergence of immune-related signaling in both regions, alongside divergent neurotransmission-related pathways in the Nacc, underscores the complexity of stress-related molecular adaptations and the precision of FKBP51-targeted interventions. Together, these results suggest that SAFit2 not only abolishes ELA-induced gene expression changes but also restores region-specific molecular functions critical for emotional regulation and social behavior.

## Discussion

In this study, we investigated if the pharmacological inhibition of the psychiatric risk factor FKBP51 can abolish the long-term behavioral and brain-wide transcriptional consequences of early life adversity. Our findings show evidence that targeting FKBP51 during or shortly after the adversity period is a viable therapeutic strategy. Using an automated, and semi-naturalistic phenotyping system, we demonstrated that ELA-induced persistent deficits in social behavior are fully restored by early-life treatment with the selective FKBP51 inhibitor SAFit2. To unravel the underlying molecular pathways related to this rescue, we performed transcriptional profiling across six stress-relevant brain regions and revealed that SAFit2 effectively normalizes ELA-induced gene expression changes, with the most pronounced effects seen in the mPFC and Nacc. Functional analysis of the rescued genes points to the reversal of dysregulated immune signaling and neuroactive ligand-receptor pathways, suggesting that FKBP51 inhibition acts on core molecular processes that underpin the embedding of early adversity. Collectively, these results establish FKBP51 as a critical pharmacological target for reversing the lasting impact of early life adversity on social behavior and brain function.

Our findings underscore previous observations, indicating that exposure to early adversity leads to a deficit in social behavior, manifested as social subordination and reduced social engagement^19,35^. This aligns with the clinical observations of social withdrawal and impaired social functioning in individuals with a history of childhood trauma^2,36^. Crucially, we demonstrate that early-life treatment with the selective FKBP51 inhibitor SAFit2 completely restored the normative social behavioral phenotype in mice. To understand the mechanism underlying this behavioral rescue, we examined the physiological hallmarks of ELA. While SAFit2 treatment did not prevent the immediate physiological stress response to the limited bedding and nesting paradigm, it did promote a faster return to baseline body weight by P25 and likely moderated the embedding of persistent dysregulation of the HPA axis. A core function of FKBP51 is that it acts as a co-chaperone of the GR-complex and suppresses the translocation of the GR-complex to the nucleus; therefore, its inhibition by SAFit2 is expected to lead to increased GR translocation and increase the efficacy of the negative feedback loop^21,37^. We propose that this pharmacological promotion of efficient glucocorticoid feedback during the critical post-trauma period conferred resilience to the stressful experience, preventing the pathological embedding of the stress response. Although SAFit2’s strong effects on the HPA axis are highlighted by the improved physiological parameters, the site of action in the offspring is currently unclear. The observed physiological effects could stem from actions restricted to the periphery (e.g., adrenal and pituitary)^38,39^, but given the compound’s brain penetrance and the observed reversal of transcriptional changes, engagement of central FKBP51 signaling is also plausible.

To uncover the molecular mechanisms underlying SAFit2-mediated rescue, we applied a pattern scoring approach designed to identify genes that are dysregulated by ELA and subsequently restored by FKBP51 inhibition. This analysis revealed a regionally diverse transcriptional landscape, with the mPFC, NAcc, and BLA exhibiting the most prominent gene expression patterning in response to ELA and its pharmacological reversal. Notably, genes such as *Fkbp5*, *Abcb1b*, and *Tlr1* showed consistent dysregulation across regions, whereas others followed region-specific trajectories, highlighting both shared and distinct molecular responses to ELA and its reversal by SAFit2. Among the consistently dysregulated genes, *Fkbp5*, a direct glucocorticoid receptor and SAFit2 target, serves as a validation marker for FKBP51 inhibition. In addition, we previously identified *Abcb1b* as a regulator of glucocorticoid secretion in the adrenal cortex, upregulated in both chronically stressed mice, and human patients with Cushing’s syndrome^40^. Our current findings extend its role to the brain, suggesting that it may act as a peripheral-central integrator of glucocorticoid dynamics. Supporting this, a common ABCB1 polymorphism (rs2032582) was also associated with a blunted cortisol response after corticotropin-releasing factor stimulation in depressed human patients. These findings align with our preclinical data and support a conserved role for ABCB1/*Abcb1b* in regulating HPA axis dynamics across species. Finally, *Tlr1*, a Toll-like receptor, has been implicated in neuroimmune signaling and psychiatric disorders^41^, also showed consistent regulation across regions. This supports the idea that innate immune pathways are integral to the transcriptional response to ELA and its reversal by FKBP51 inhibition.

These results also build on our previous findings^19^, which showed that ELA induces distinct transcriptional signatures in GABAergic and glutamatergic neurons of the ventral hippocampus. Using a comparable pattern scoring framework, we identified transcriptional motifs (*i.e.*, chronic, primed, inverted, and blunted) that varied depending on ELA exposure, and a second hit by acute stress, revealing nuanced gene regulation dynamics. Our new bulk RNA-seq findings extend this concept, showing that transcriptional patterning is not confined to the vHPC, but instead reflects a broader, regionally distributed molecular signature of ELA that is responsive to FKBP51 inhibition. Importantly, the convergence of gene patterning across regions, particularly within immune and neuroactive ligand-receptor pathways, parallels the functional clusters reported by Kos et al^19^. Together, these findings suggest that FKBP51 inhibition with SAFit2 not only rescues behavioral impairments but also re-establishes key molecular programs disrupted by early adversity. This convergence reinforces FKBP51’s role as a key regulator of the transcriptional embedding of ELA and a promising target for its reversal.

## Conclusion

Taken together, our data show that early-life inhibition of FKBP51 with SAFit2 prevents the embedding of early life adversity into persistent social subordination and brain-wide transcriptional dysregulation. The administration of SAFit2 during the early life adversity window restored social behavior and social hierarchy, while normalizing ELA-driven gene expression changes across several stress-related brain regions, with pronounced effects in mPFC and Nacc. The rescued transcriptomic signatures converged on immunoregulatory processes and neuroactive ligand–receptor signaling, consistent with a SAFit2-mediated strengthening of glucocorticoid feedback. This positions FKBP51 as a critical pharmacological target for reversing the lasting impact of early life adversity on social behavior and brain function, offering a path toward developing targeted, preventative treatments for ELA-related psychiatric disorders.

## Acknowledgements

The authors thank Prof. Gerhard Winter and Katharina Kopp from the Department of Pharmacy at the LMU Munich, Germany for the production of the vesicular phospholipid gels. Furthermore, the authors thank Bianca Schmid and Rainer Stoffel for their technical assistance, and the DeepLabCut development team for creating and maintaining the DeepLabCut software. We acknowledge the technical support of the Core Facility Genomics at Helmholtz Zentrum München. The computations/data handling was enabled by resources provided by the National Academic Infrastructure for Supercomputing in Sweden (NAISS), partially funded by the Swedish Research Council through grant agreement no. 2022-06725.

## Author contributions

**Conceptualization**: J.B., J.P.L., M.V.S; **Experimental design**: J.B., X.J., J.P.L., M.V.S.; **Data analysis**: J.B., X.J., C.N.; **Software**: X.J., S.G., C.S.C.; **Investigation**: J.B., X.J., D.H., C.F., C.N.; **Validation**: J.B., X.J.; **Resources**: M.U., G.W., F.H., A.C., J.P.L., M.V.S.; **Data Curation**: J.B., X.J., P.F., J.P.L., M.V.S.; **Visualization**: J.B., X.J.; **Supervision**: P.F., J.P.L., M.V.S.; **Writing – Original Draft**: J.B., X.J., J.P.L., M.V.S.; **Writing – Editing**: All authors; **Project Administration**: J.P.L., M.V.S.; **Funding Acquisition**: J.P.L., M.V.S.

## Data and Materials Availability

All data needed to evaluate the conclusions in the paper are present in the paper and/or the Supplementary Materials. Additional data related to this paper may be requested from the authors. The transcriptomics data sets and processed data files are available upon request before publication and will be made publicly available upon publication.

## Methods

### Animals and housing conditions

Wild-type adult male and female CD1 mice (2-3 months old) from the in-house breeding facility of the Max Planck Institute of Psychiatry were used as breeders (F0). Their male offspring (F1) served as experimental animals. At weaning, F1 mice were housed in groups of four, with each cage containing animals from different litters, depending on the condition (see experimental design section). Mice were kept in individually ventilated cages (IVC; 30 × 16 × 16 cm, Tecniplast Green Line - GM500) connected to a central airflow system, under standard housing conditions: 12 h/12 h light–dark cycle (lights on at 7:00 a.m.), ambient temperature 23 ± 1 °C, and relative humidity 55%. Food (Altromin 1324, Altromin GmbH, Germany) and water were available *ad libitum*. All experimental procedures were approved by the Committee for the Care and Use of Laboratory Animals of the Government of Upper Bavaria, Germany, and were conducted in accordance with the European Communities Council Directive 2010/63/EU.

### Experimental design

At postnatal day 2 (P02), litters were randomly assigned to control (Ctrl) or early life adversity (ELA) conditions, and dams were treated with either the FKBP51 inhibitor SAFit2 or vehicle (Veh). At weaning (P25), offspring were housed according to a 1:2:1 design (Ctrl_Veh:ELA_Veh:ELA_SAFit2). This design resulted in cages containing two stress-affected animals (ELA_Veh), one non-affected control (Ctrl_Veh), and one potentially stress-resilient animal (ELA_SAFit2).

### The early life adversity paradigm: limited bedding and nesting

Early life stress (ELA) was performed using the limited bedding and nesting (LBN) paradigm to induce chronic stress towards the mother and pups during P02 to P09, as previously described by Rice et al^30^. At P02, all litters were transferred to new IVCs and randomly assigned to the non-stressed (NS) or stressed (ELA) condition. If necessary, the litters were culled to a maximum of 10 animals per litter. The ELA litters were placed on a stainless-steel mesh (McNichols) and were provided with limited nesting material (1/2 square of Nestlets, Indulab). The NS animals were placed in an IVC with a standard amount of bedding material and were provided with a sufficient amount of nesting material (2 squares of Nestlets). All litters were left undisturbed until P09, after which they returned to standard housing conditions. The pups were weaned in same-sex groups with a maximum of four animals per cage.

### Pharmacological treatment with SAFit2

At P02, coinciding with the onset of the ELA paradigm, dams received a single subcutaneous injection of SAFit2 vesicular phospholipid lipid gel^42^ (VPG; 200 µl, containing 10 mg SAFit2 per gram of gel). This formulation provides sustained release, resulting in stable plasma and brain concentrations with a half-life of approximately 7 days^23,43^. Pups were exposed to SAFit2 via maternal milk (Figure 1). Control animals received an injection of empty VPG without SAFit2 (Veh).

### The measurement of adrenals and corticosterone at P09 and adulthood

One week after the behavioral tests, adult animals were weighed and sacrificed by decapitation. Trunk blood was collected in EDTA-coated microcentrifuge tubes (Kabe Labortechnik, Germany) and immediately placed on ice. Samples were centrifuged at 4 °C for 15 min at 8,000 rpm, after which plasma was separated and stored at −80 °C until further analysis.

A separate cohort of mice was used to obtain corticosterone (CORT) measurements directly after ELA at P09. On the morning of P09, litters were kept in their home cage. The dam was sacrificed first, followed by the pups, which were kept in the nest until sacrifice to minimize additional acute stress. Trunk blood was collected and processed as described for adult samples. Plasma CORT levels were measured in duplicate using radioimmunoassay, following the manufacturer’s protocol (MP Biomedicals, Eschwege, Germany).

Adrenal glands were dissected at P09 and P70 (adulthood) and kept in 0.9% NaCl at 4 °C until further processing, which included removal of surrounding fat tissue and weighing. Relative adrenal weight was calculated as the ratio of combined adrenal weight (left and right) to body weight measured prior to sacrifice.

### Measurement of SAFit2 concentration

Mice blood plasma samples were analysed using the combined high-performance liquid chromatography/mass spectrometry (HPLC/MS-MS) technique. Analysis was performed using a Shimadzu Nexera X2 (Shimadzu, Duisburg, Germany) liquid chromatograph which was interfaced to the ESI source of a Sciex QTrap 5500 (Sciex, Darmstadt, Germany) triple quadrupole mass spectrometer. All samples were prepared using Ostro protein precipitation and phospholipid removal plates (Waters, Eschborn, Germany).

Chromatography was accomplished using a gradient elution in a Accucore RP-MS column (100 x 2,1mm, 2,6µm Thermo Scientific, Dreieich, Germany) at a flow rate of 0.5 ml/min and 30 °C. The composition of eluent B was methanol with 10mM ammonium formate with 0,1% formic acid and water with 10mM ammonium formate with 0,1% formic acid as eluent A. The gradient was 0-3 min 60% B, 3-4,5 min 60-90% B, 0,5 min held at 90% B, 5-5,2 min 90-60% B and 5,2-6 min 60% B. The total run time was 6 min and the injection volume was 2µl. The ion source was operated in the positive mode at 500°C, and multiple reaction monitoring (MRM) collision-induced dissociation (CID) were performed using nitrogen gas as the collision gas. Deuterated SAFit2 (SAFit2-D3) was used as internal standard. The retention time and transitions monitored during analysis for the analytes were as shown in the following table.

| Compound | Used as | Q1_Mass | Q3_Mass | RT [min] | DP [V] | EP [V] | CE [V] | CXP [V] |
| --- | --- | --- | --- | --- | --- | --- | --- | --- |
| SAFit2 | Quantifier | 803,2 | 384,2 | 4,95 | 141 | 10 | 41 | 18 |
| SAFit2 | Qualifier | 803,2 | 114,2 | 4,95 | 141 | 10 | 67 | 8 |
| SAFit2-D3 | Internal Standard | 806,4 | 384,3 | 4,95 | 106 | 10 | 41 | 26 |

### Social Box (SB) behavioral assessment

Groups of male non-sibling CD-1 mice consisting of one Ctrl_Veh, two ELA_Veh and one ELA_SAFit2 animals were housed together from weaning. The fur of the animals was painted in different colors to keep track of their identity. Behavioral assessment in SBs took place at postnatal day 30 and 61, corresponding to adolescence and young adulthood. The use of SB phenotyping systems was described in detail previously ^33,44^. Each group was housed in a 60 cm x 60 cm living environment with two feeders for water and food, an enclosed nest and a smaller open nest, an s-wall, and two ramps. The arenas were illuminated with around 200 lux during the light phase (12 hours) and 2 lux during the dark phase (12 hours). For each assessment, recording of the animals was done continuously in four dark phases and three light phases with a color-sensitive camera.

A custom neural network was trained with Deeplabcut^45,46^ (version 2.3.5) to analyze the videos and track the positions of body parts. The tracking data were then parsed into ethologically relevant behavioral features, including chasing, sniffing, David Score (DS), and interaction with SB objects, using a combination of machine-learning based^47^ tools and heuristic algorithms. Behavioral data were binned into multiple time windows (1-, 2-, 4-, 6-, 12-hour bins) to capture both short-term and long-term dynamics. The duration and event count of each behavior were normalized by the time each animal spent outside the nest.

### Analysis of behavior readouts

The influence of ELA and SAFit2 treatment on the behavioral features were assessed using linear mixed-effects models implemented in the “nlme” package^48^ in R (version 4.4.0). For each feature, values were normalized and log-transformed prior to analysis. The model included condition, day and time window as fixed effects, and individual mice nested within groups as a random effect to account for repeated measurements. To visualize the changes and significance of each feature in ELA_Veh and ELA_SAFit2 groups compared to Ctrl_Veh group, the coefficients and p-values for all features were combined, and false discovery rate (FDR) adjustment was applied on the p-values. The analyses were stratified by smaller time windows for the normalized David’s score and specific behaviors, and two-way repeated-measures ANOVA was performed to assess condition-specific effects over time. Principal component analysis (PCA) was performed on normalized behavioral features in different time windows and nights using the “scikit-learn” implementation in Python. The first two principal components (PC1 and PC2) were extracted for visualization and downstream statistical analysis.

### Transcriptomics

Following SB phenotyping under basal conditions, animals were sacrificed and brains were rapidly removed, snap-frozen in 2-methylbutane cooled on dry ice, and stored at −80 °C until further processing. Coronal sections (250 µm) were cut on a cryostat, and tissue punches were collected from the following regions (anteroposterior coordinates relative to bregma): medial prefrontal cortex, infralimbic and prelimbic (mPFC; AP +2.10 to +1.54 mm); anterior cingulate cortex (ACC; AP +2.10 to +0.50 mm); nucleus accumbens shell (Nacc; AP +1.94 to +0.98 mm); basolateral amygdala (BLA; AP −0.70 to −1.91 mm); dorsomedial thalamus (DMT; AP −0.70 to −1.91 mm); and ventral hippocampus (vHipp; AP −2.92 to −3.64 mm). Punches were stored at −80 °C until RNA isolation. Total RNA was isolated from punches using the miRNeasy Mini Kit (QIAGEN, Venlo, NL; cat. no. 1038703) according to the manufacturer’s instructions, using RNase-free technique throughout.

After RNA isolation, RNA integrity number (RIN) was measured using the Agilent 2100 Bioanalyzer system and quantified by Qubit (Thermo Fisher Scientific). RNAs with a RIN value > 7 were selected for mRNA sequencing (poly-A selected). The libraries were prepared using the Illumina Stranded mRNA Prep, Ligation kit (Illumina), following the kit’s instructions. After a final QC, the libraries were sequenced in a paired-end mode (2×100 bases) in the Novaseq6000 sequencer (Illumina) using an S4.200 flowcell with a depth of ≥30 million paired-reads per sample.

### Quality control and differential expression analysis

The raw counts matrix was processed in R (version 4.4.0). Lowly expressed and non-informative genes were filtered prior to analysis. Specifically, genes with low cumulative count (<100) and genes expressed in few samples (<6) were discarded. Additionally, non-protein-coding genes (according to Ensembl gene type annotation) were also discarded. The raw counts of samples from the same brain region were grouped and filtered together, and the filtered counts for each brain region were used for the subsequent differential expression analyses. Differential gene expression analysis was performed in R (version 4.4.0) using the “DESeq2” package^50^. Pairwise comparisons were made within the three conditions: ELA_Veh versus Ctrl_Veh, ELA_SAFit2 versus ELA_Veh and ELA_SAFit2 versus Ctrl_Veh. Genes with |log2FC| > 0.5 in ELA_Veh versus Control_Veh or ELA_SAFit2 versus ELA_Veh were filtered for further analysis.

### Transcriptional patterning calculations

To identify genes with expression changes that are induced by ELA and restored by SAFit2, a gene patterning score is calculated based of the log2FC of each gene in the three comparisons:

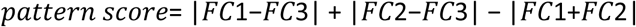

Where:

FC1 = log2FC ELA_Veh versus Ctrl_Veh

FC2 = log2FC ELA_SAFit2 versus ELA_Veh

FC3 = log2FC ELA_SAFit2 versus Ctrl_Veh.

A higher gene patterning score suggests an increased likelihood for the expression pattern to be affected by both ELA and restored by SAFit2 treatment. The distributions of gene patterning scores were visualized and compared across different brain regions. The effect of brain regions was assessed with one-way ANOVA following post hoc estimated marginal means with effect contrasts (difference from the grand mean) using the “emmeans” package^49^ in R (version 4.4.0). The log2FC of the top 30 highest scoring genes in each region were visualized as a heatmap using “ggplot2” in R (version 4.4.0).

### Metascape gene network analysis

The metascape gene annotation and functional enrichment tool^34^ was used to analyze genes with high patterning scores (above the 25th percentile) from mPFC and Nacc. The genes were divided into three lists, which consist of genes showing patterning only in mPFC, genes showing patterning only in Nacc, and genes showing patterning in both. The three gene lists were uploaded using the “multiple gene list” setting and analyzed with the express analysis settings. The gene enrichment clusters were visualized together with the contribution of the three gene sets (mPFC specific, Nacc specific, overlapping) to each of the identified clusters.

### Statistics

Statistical analyses and data visualization for the physiological hallmarks of early life stress (Figure 1) were performed using GraphPad Prism 10 and RStudio (R version 4.4.0). Outliers were identified with Grubbs’ test and excluded when significant. Assumptions of normality were evaluated with the Anderson-Darling test. When assumptions were violated, non-parametric alternatives were applied. Comparisons among three groups were conducted using one-way ANOVA (parametric) or Kruskal-Wallis test (non-parametric). Significant main effects were followed by post hoc analyses using the Holm–Šidák test (parametric) or Dunn’s test (non-parametric). Bar graphs are presented as the mean ± standard error of the mean (SEM). Data were considered significant at p<0.05 (*), and further significance was represented as p<0.01 (**), p<0.001 (***), and p<0.0001 (****).

## Supplemental figures

**Supplementary Figure 1.**
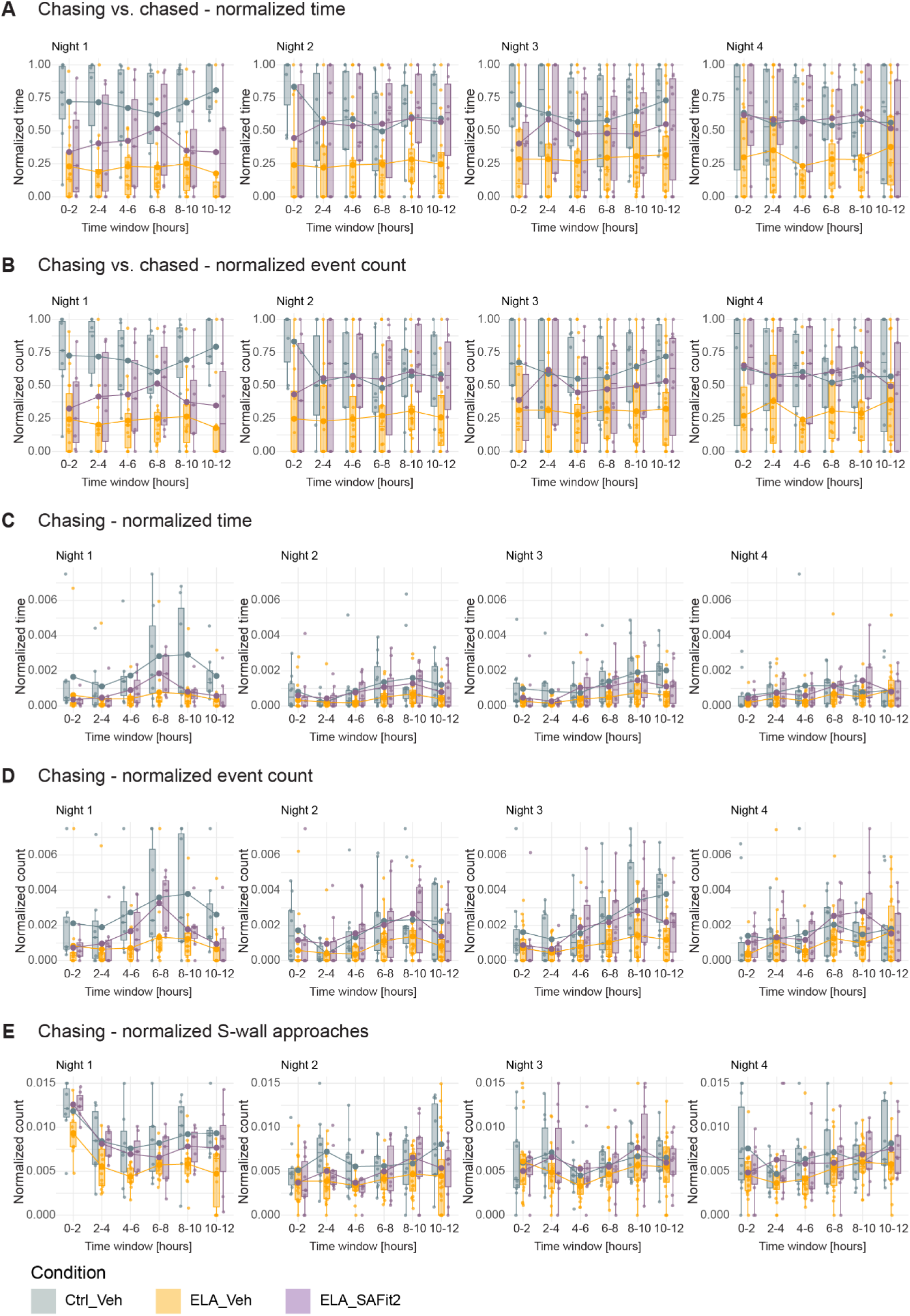
Time-resolved profiles of significant behavioral features. A-D). Behavioral readouts from the active phases were segmented into 2-hour time windows and compared across Ctrl_Veh, ELA_Veh and ELA_SAFit2 conditions. **A).** Ratio between the time spent chasing and the time being chased. **B).** Ratio between the event count of chasing and being chased. **C).** The total time spent chasing. **D).** The event count of chasing. **E).** The number of approaches to the s-wall. The time and number of events are normalized by the total time the animal spent outside the nest.

**Supplementary Figure 2.**
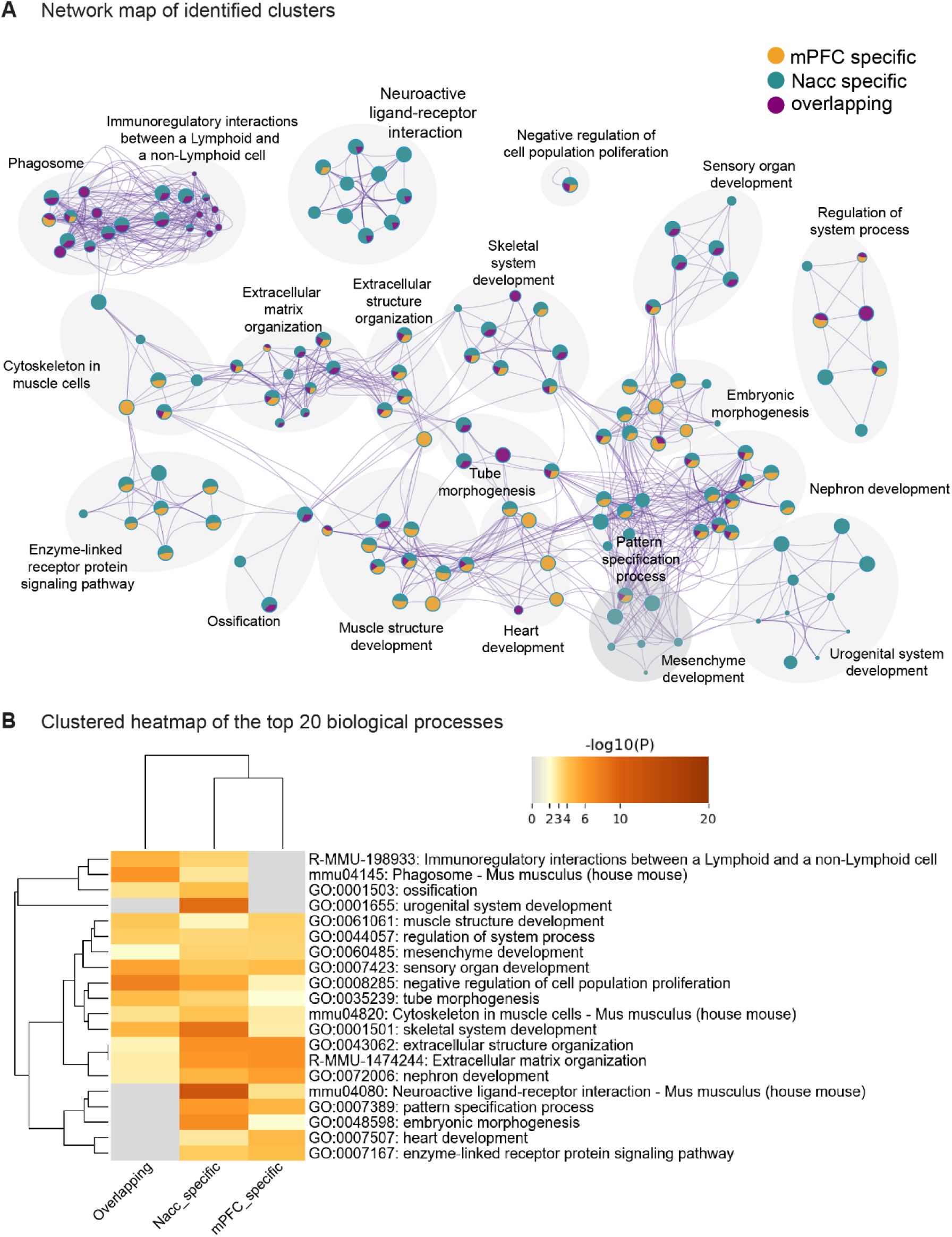
Functional clustering of genes with elevated pattern scores in the mPFC and Nacc. **A).** Network map visualization of the identified functional clusters. The pathways are colored with mPFC-specific, Nacc-specific and overlapping (present in both regions). **B).** Clustered heatmap of the top 20 biological processes enriched in mPFC-specific, Nacc-specific and overlapping genes.

